# A Multi-layer, Self-aligning Hydrogel Micro-molding Process Offering a Fabrication Route to Perfusable 3D *In-Vitro* Microvasculature

**DOI:** 10.1101/242156

**Authors:** Hossein Heidari, Hayden Taylor

## Abstract

The *in-vitro* fabrication of hierarchical biological systems such as human vasculature, which are made up of two or more cell types with intricate co-culture architectures, is by far one of the most complicated challenges that tissue engineers have faced. Here, we introduce a versatile method to create multi-layered, cell-laden hydrogel microstructures with coaxial geometries and heterogeneous mechanical and biological properties. The technique can be used to build *in-vitro* vascular networks that are fully embedded in hydrogels of physiologically realistic mechanical stiffness. Our technique produces free-standing 3D structures, eliminating rigid polymeric surfaces from the vicinity of cells and allowing layers of multiple cell types to be defined with tailored extracellular matrix (ECM) composition and stiffness, and in direct contact with each other. We demonstrate co-axial geometries with diameters ranging from 200–2000 μm and layer thicknesses as small as 50–200 µm in agarose– collagen (AC) composite hydrogels. Coaxial geometries with such fine feature sizes are beyond the capabilities of most bioprinting techniques. A potential application of such a structure is to simulate vascular networks in the brain with endothelial cells surrounded by multiple layers of pericytes and other glial cells. For this purpose, the composition and mechanical properties of the composite AC hydrogels have been optimized for cell viability and biological performance of endothelial and glial cell types in both 2D and 3D culture modes. Multi-layered vascular constructs with an endothelial layer surrounded by layers of glial cells have been fabricated. This prototype *in-vitro* model resembles vascular geometries and opens the way for complex multi-luminal blood vessels to be fabricated.

## 1. Introduction

The evolution of tissue engineering over the past few decades can be attributed to the advent of engineered biomaterials, novel biofabrication techniques, and powerful microfluidic platforms. Applications of tissue engineering are found throughout regenerative medicine, drug discovery and screening, fundamental biological studies using *in-vitro* disease models, and *in-vitro* meat production (*e.g.* [1]). In all of these areas, however, it has not yet proved possible fully to bridge the gap between small ‘on-chip’ models and large ‘transplantable’ ones. In other words, organ-scale bio-printed constructs lack full biological function and performance, and microfluidic and on-chip models, however accurate, are not easily scalable to the dimensions of a transplantable organ. Hence, when it comes to producing dense volumetric samples of tissue, thicknesses even on the millimeter scale are far from reach. An example of such a challenge lies in the production of lab-grown meat, where enormous effort was required even to produce a single centimeter-thick burger with almost 20,000 millimetric strands of muscle cell bundles [1].

In the present work, we focus on human microvasculature for two reasons. Firstly, one of the main reasons that we do not have such bulk tissue models is the lack of biologically relevant 3D perfusable vascular networks to facilitate transport of species to and from the cells. Secondly, one of the most important systems that is involved in almost all disease models is the human vasculature, and as we will briefly discuss, most of the available *in-vitro* disease models lack the geometrical complexity as well as the multitude of vasculature-specific cell types. As we know, every category of human blood vessel, including coronaries, veins, and cerebral vessels, is characterized by specific cell types, tissue layer thicknesses and lumen diameters that result in a unique set of values for permeability, compliance and integrity of that exchange site. Cerebral capillaries are characterized by a very thin and tightly packed layer of endothelial cells covered by pericytes and glial cells, whereas coronary arteries consist of a thick layer of smooth muscle cells and elastic tissue offering huge compliance and much less integrity. Therefore, in order to satisfy the biological requirements of a particular category of blood vessel, an *in-vitro* model should also include the same multilayered tissue composition, mechanical properties and architecture.

A very common approach to *in-vitro* modelling of vasculature is producing a two-dimensional (2D) interface that captures the specificity of the vessel walls and could be utilized for most transport-related studies and disease models. In the past years, 2D models of vasculature such as the artificial membrane model [2], cone and plate model [3] and hydrogel-based 2D models [4] have evolved into a standard trans-well membrane system which has been widely used due to its ease of use and convenience [5], [6]. More recently, this classic trans-well system has been further upgraded to an on-chip platform with integrated microfluidics, which has been used to construct *in-vitro* blood–brain barrier (BBB) models such as the ‘BBB-on-chip’ [7] and the ‘micro-BBB’ [8].

Falling in the same category of 2D interfaces are microfluidic chips that utilize diffusion chambers injected with extra-cellular matrix (ECM) that separate multiple culture channels. These models rely on the diffusion of agents through the channel–ECM interfaces and through the ECM regions themselves [9], [10]. Although these devices enable controlled diffusion of cell-secreted factors and chemicals, they may suppress important signaling pathways such as intercellular mechano-transduction because neighboring cell types are kept isolated.

Perhaps more importantly, in all of the models falling into this category, endothelia are exposed to materials that are far stiffer than they encounter in the body. For example, microfluidic devices, even if injected with ECM, frequently incorporate elastomeric surfaces that are 100–1000 times stiffer than typical ECM; meanwhile, the polystyrene trans-well membranes that we mentioned earlier are around 10^6^ times higher in elastic modulus. Moreover, increasing the number of vessel wall layers above two is exceptionally difficult with any membrane-based system. The compliance of the membrane in response to the actual physical flow conditions and pressure is dominated by the supporting substrate stiffness rather than by the actual tissue.

A second approach to *in-vitro* modelling of vasculature is inspired by vasculogenesis and angiogenesis and involves the natural growth of endothelial sprouts within a 3D matrix. Microvascular networks-on-a-chip fall into this category, and many groups have successfully produced perfusable endothelialized sprouts that extend within a gel chamber [11]–[13]. This method offers natural network geometries and vascular growth. Nevertheless, there are a few significant issues when it comes to practicality in the aforementioned main target areas of organ transplants and disease models. Firstly, the natural lumen formation and vascularization enables only a single-layered lumen with only one cell type. It is impossible then to produce a secondary layer of cells surrounding those sprouts; in the best case one could add a second cell type by embedding cells within the surrounding ECM. Therefore, this approach is effectively limited to single-layered vessels. Furthermore, the diameter and thickness of each lumen as well as the inlets and outlets of the channels are very difficult to control. These limitations are especially undesirable for most *in-vitro* disease models and drug permeability studies, since they further complicate the experimentation and monitoring setup.

Finally, we have an emerging class of novel fabrication methods such as sacrificial molding [14], UV-based bio-printing in which materials are photocrosslinked [15], [16], nozzle-based 3D bio-printing [17], and freeform embedding of hydrogels within a sacrificial gel (‘FRESH’) [18]. These methods deliver much higher control over the spatial patterning of tissue than the previous two classes of method, and in addition are able to print multiple cell types. Even these methods, though, have their limitations. Sacrificial molding suffers from the same complications associated with seeding multiple cell types and producing multiple concentric shells and layers, since it is only possible to remove the sacrificial core once. Furthermore, removal of the sacrificial core can be a very challenging task at capillary-scale diameters. UV-based bio-printing is very similar to stereolithography and is therefore limited to printing with only a single material; the gel composition and cell type cannot vary throughout the printing process. Nozzle-based printing and FRESH have been used to produce approximations to a cylindrical vessel with a resolution of a few hundred microns [17], [18]. However, issues associated with nozzle positioning and tissue layer alignment are expected to be pronounced if one were to attempt to scale down these processes to tens of micrometers. It should be noted that the advent of nozzles that can print hollow cylindrical tubes of hydrogel does potentially add a lot of capability to such extrusion-based 3D bioprinters [19]. Nevertheless, scaling such extrusion-based methods for hollow fibers below 100 µm can introduce many technical difficulties that have not yet been overcome. Moreover, controlling the thickness as well as the stiffness of each layer is not currently possible using this class of vasculature models.

We believe that architecture, geometry and topology of the microvascular structure are extremely important, not only from the biological perspectives briefly explained above but also from a fluid mechanics perspective, where the critical effects of biological flow such as pulsatility and wall compliance, and the induced wall shear stress [20], [21] can significantly alter cell growth, morphology and arrangement as well as mass transport either throughout the culture process or after the *in-vitro* model is prepared and ready to experiment with. Here, we present a versatile and accurate yet extremely convenient and cost-efficient fabrication route that can accurately capture the complex multi-cellular composition, cellular interactions and vascular architecture of human vasculature to a greater level than most of the aforementioned techniques.

## 2. Materials and methods

### 2.1. Fabrication of the 3D structures; multilayer micro-molding

In order to achieve the multi-layer coaxial structures, our process uses a sequence of molded thermal gelation steps to build the microstructure (Fig 1). A 3D design of the vasculature is obtained using either a custom-designed vascular network model generated with computer-aided design (CAD) software, or, in principle, a standard triangle language (STL) representation of a computed tomography (CT)-based z-stack image of *in-vivo* human vasculature. For each layer of the coaxial structure desired, the model is then converted into a corresponding, separable tw o-part mold with the required alignment p rotrusions and other fixtures. The molds are 3-printed using an Objet Connex 260 3D inkjet printer with a ominal layer thicknes s of 16 µm. The printed resin material is VeroCle ar™, an acrylate-based photocurable material. Other mol d-making p rocesses wi th higher resolution could also be used.

**Figure 1.**
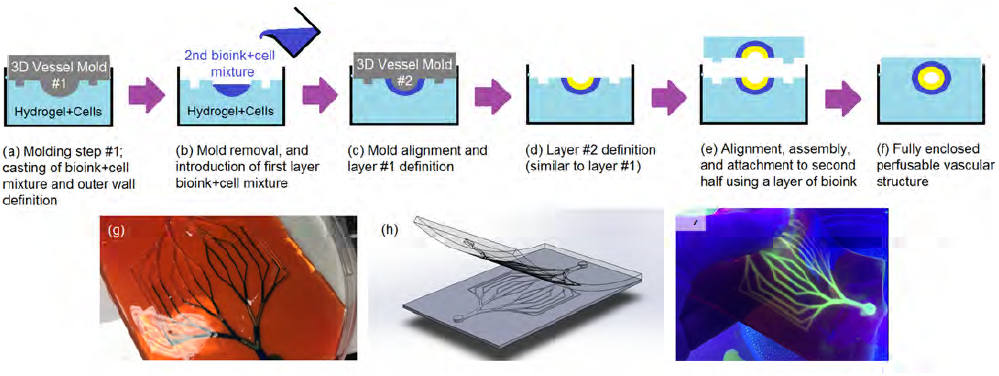
Fabrication method: (a–f) process flow; (g) flow in the assembled perfusable network; (h) illustration of 3D-printed mo ld design to p roduce that ne twork; (i) flexibility and compliance of the molded network.

In order to achieve multiple layers in t he hydrogel structures, one molding step is followed by another with the same layout but smaller-dimeter protrusions. The material to be molded is heated,and starting with the mold wit the larges t diameter of protrusions, the outermost surface of the lumens is defined by casting the hydrogel–cell mixture representing the surrounding tissue (Fig 1a).After each casting step, the molded material is allowed to gel thermally as it cools to below the gelation temperature. Subsequetly, anothe mold with smaller-diameter protrusions is used and a half-cylindrical shel of hydrog l–cell mixture is cast into the intermediate space (Fig 1b–c). This sequence can be exte nded to multiple layers at different radial locations (*e.g.*Fig 1d). Layer-to-layer registration at each step is enabled by the self-centering action of mold protrusions within previously molded hydrogel features. Posts printed onto the molds further enhance registration, and enable the two halves of the hydrogel construct to interlock and form fully enclosed, coaxial structures (Fig 1e– f).

Following the above process, for each new hydrogel layer that is deposited we can culture cells either in the form of a 2D monolayer on the surface of the solidified hydrogel, or as a 3D cell population that is mixed into the material prior to casting so that it becomes uniformly embedded within the hydrogel layer. Therefore, our fabrication process combines 2D and 3D culture modalities into a multi-layered model.

### 2.2. Composite agarose–collagen hydrogels

Agarose is a thermosensitive natural polysaccharide that consists of copolymers of 1,4-linked 3,6-anhydro-α-L-galactose and 1,3-linked β-galactose. Since the gelation mechanism of agarose is through hydrogen bonding and chain entanglement, the gelation temperature may be configured based on the desired application in a range between 17 °C and 40 °C, depending on the degree of hydroxylethyl substitution [22]. Interestingly, the melting temperature of agarose is approximately 40 °C greater than the gelation temperature, displaying a special hysteresis behavior. This property is especially desirable for our application as it enables rapid polymerization and irreversible gelation at the proximity of incubation temperatures, therefore eliminating swelling and post-fabrication deformations in our temperature range of interest.

As will be explained, agarose-based hydrogels show exceptional mechanical performance, and are moldable even at very low polymeric concentrations (1% w/v and lower) and stiffness values (1– 10 kPa and less). Unfortunately, despite their desirable mechanical properties, prior use of agarose-based hydrogels as an *in-vitro* ECM has been limited. We will show that this limitation could be mainly attributed to their lack of cell-adhesive domains and relatively low cell viability in long-term cultures. One way of solving this biocompatibility issue is by immobilizing adhesion ligands on the crosslinked chains through the activation of hydroxyl groups [23]. An alternative way is the use of binary composites of agarose and other natural biopolymers and ECM proteins [5]. The latter solution offers more degrees of freedom and multi-parameter tunability, via the independent modulation of each component of the hydrogel. Here, we have studied a wide range of binary agarose–collagen (AC) hydrogels for their mechanical performance, manufacturability and biocompatibility in the two main culture modalities that are incorporated in vascular architectures:(1) 2D culture for endothelial lumen surfaces, and (2) 3D culture for the surrounding tissue.

Here, we have studied a wide range of AC hydrogels with varying concentrations (w/v) of agarose (0.25, 0.5, 1.0 and 2.0%), all of which can be regarded as very low polymeric content gels. The concentration (w/v) of the second component (collagen) was also modulated (0.025, 0.05, 0.1 and 0.2%). In order to prepare the blends, based on the desired final concentrations, each component was prepared individually, and in all cases cells were mixed with the collagen component first and then the agarose solution was added to the mixture. In the case of agarose, high-melting-point agarose powder (J.T. Baker, Center Valley, PA, USA) with a gelation temperature of 36 °C was dissolved in sterile Dulbecco’s Phosphate Buffered Saline (DPBS) (Gibco, Rockville, MD, USA) and the mixture was heated in a microwave oven for two consecutive time periods of 30 seconds each, in order to avoid boiling of the buffer, until the solution became fully transparent. The agarose solution was then filter-sterilized using a 0.22 µm pore-size syringe filter, and was kept at 45 °C in a controlled environment until needed. For the case of collagen, type 1 rat tail collagen in solution format (Corning, Corning, NY, USA) at a concentration of a 4.1 mg/mL was diluted in tissue culture grade deionized water to target concentrations.

### 2.3. Cell culture procedure

The HMEC cell line (ATCC CRL-3243, American Type Culture Collection) is a microvascular endothelium cell line with endothelial morphology. The HMEC cells were cultured in MCDB 131 (Thermo-Fisher Scientific, Waltham, MA, USA) supplemented with 10% fetal bovine serum, 10 ng/mL endothelial growth factor (EGF) (Thermo-Fisher Scientific), and 1% penicillin/streptomycin at 37 °C in a humidified 5% CO2 atmosphere. The SH-SY5Y cell line (ATCC CRL-2266, American Type Culture Collection) is a neuroblastoma cell line with epithelial morphology. The SY5Y cells were cultured in Dulbecco’s Modified Eagle’s Medium (Gibco) supplemented with 10% fetal bovine serum (Gibco), 1% NEAA, 1% sodium pyruvate and 1% penicillin/streptomycin (Gibco) at 37 °C in a humidified 5% CO2 atmosphere. All subcultures were passaged at 80–90% confluence and throughout the experiments, doubling time was almost 48 and 32 hours for the SY5Y and HMEC cell lines respectively. The cells were recovered at this stage after detachment from the culture plates using fresh 0.05% trypsin-EDTA (1X) solution (Gibco) and then either passaged, or used in the experiments. In all long-term culture experiments, the medium was refreshed every four days.

### 2.4. Cell-laden hydrogel for 3D culture

For the 3D culture mode, due to the rapid cooling and thermal crosslinking of agarose, the previously described collagen solution was initially mixed with the media solution containing cells at a target density. The agarose solution was then allowed to cool to below 40 °C, after which it was pipetted into the mixture. The blend was mixed thoroughly, after which it was immediately transferred and cast into the final container, where it was stamped with the respective mold. After a few minutes and the completion of the thermal gelation process at room temperature, the samples were covered with media, or, in the case of multiple-layer structures, the casting–molding process was repeated with another layer of the cell–hydrogel mixture.

### 2.5. Hydrogel surface coatings for 2D culture

In order to achieve an epithelial or endothelial lumen, we had to engineer the cell adhesion properties of the agarose surface to enable a 2D culture mode. Unlike many hydrogels such as collagen, gelatin, and methacrylates of polyethylene glycol, gelatin and hyaluronic acid, that have been studied intensively for surface enhancements, agarose hydrogels have not been explored as much. We therefore conducted a study of various surface coatings that would enable cell adhesion and spreading on agarose surfaces. The presented sub-study is critical to the determination of the potential of this hydrogel for the proposed model.

The main surface coatings that we have studied are poly-d-lysine (PDL), Matrigel, collagen, and human-derived fibronectin (hFN). Unfortunately, previously reported preparation and coating protocols, and the recommended concentrations of these coatings, have mostly been optimized for the rigid plastic surfaces of commercially available culture dishes. The applicability of such protocols to hydrogel surfaces has not been systematically investigated in the literature. Through a set of experiments, we came up with our own optimal procedures and here, we shall briefly describe the coating process configurations that have been used for each material.

PDL hydrobromide in powder format (Sigma-Aldrich, St. Louis, MO, USA) was dissolved in sterile DPBS to a final concentration of 0.1 mg/mL and this stock solution was then diluted over a range of concentrations (0.01, 0.025, 0.05, 0.1, and 0.25 mg/mL) in order to find the optimal coating concentration. The hydrogel substrates were covered with 250 μL/cm^2^ of the diluted PDL solutions. The samples were then incubated at 37 °C and 5% CO2 for an hour. They were then removed from the incubator and washed twice with sterile DPBS, after which they became ready for cell seeding or the application of the secondary coating.

Matrigel in solution format (Corning) was stored at –20 °C, thawed overnight at 2–8 °C and then kept over ice throughout the dilution process. Pipette tips were also kept at 2–8 °C before coming in contact with the Matrigel. The Matrigel was diluted in sterile DPBS to a final concentration of 1 mg/mL which was then kept as a stock solution and further diluted to a range of concentrations (0.1, 0.2, and 0.4 mg/mL). The hydrogel substrates were then covered with 50 µL/cm^2^ of the diluted solutions and incubated for half an hour, after which the residuals were washed away with DPBS.

Collagen type 1 stock solution with a concentration of 4.1 mg/mL was diluted in sterile tissue grade deionized water over a range of concentrations (0.5, 1.0, and 2.0 mg/mL) and then applied to the hydrogel surfaces at 200 μL/cm^2^. The hFN in powder format (Corning) was dissolved to 1 mg/mL in sterile tissue grade deionized water, frozen and stored at –20 °C. The solution was then further diluted to 50 μg/mL and used to coat the surface of the hydrogel substrates at 100 μL/cm^2^. The coated substrates were then covered with media containing HMEC and SY5Y cells at concentrations of 8.4 × 10^5^ and 13.4 × 10^5^ cells/μL respectively.

### 2.6. Mechanical characterization

Different experimental tools and methods such as rheological characterization, extensometry and nano-indentation have been used to assess the mechanical performance of hydrogels, synthetic ECM scaffolds and bio-polymers. Here we are interested in analyzing stiffness of the gels *i.e.* Young’s modulus or the elastic component of the viscoelastic behavior of our hydrogels. Although rheological data can also be translated to elasticity via the incorporation of an equivalent spring and damper model, the characterization method still falls into the bulk category and despite the breadth of information that this method offers, it is still unable to evaluate accurately what cells are actually sensing at the microscopic scale. Nano-indentation and atomic force microscopy (AFM) are among the most accurate characterization tools in this sense. Between the two of these techniques, nano-indentation was chosen because bio-AFM systems still suffer from inaccuracies associated with surface tension of the cantilever probe with the wet hydrogel samples. In the case of indentation, the indenter fully deforms the sample and loading/unloading data used to calculate the elastic modulus are collected at the extremum of the compression profile, therefore eliminating any errors associated with the weak tip–sample interaction domain.

A Hysitron TI-950 Triboindenter nano-indentation system (Hysitron, Eden Prairie, MN, USA) was used to evaluate the modulus of elasticity of the samples. Hydrogel samples were prepared inside custom-made 3 mm-tall cubic containers and a 1 mm layer of deionized water covered the top surface of the samples in order to keep them hydrated throughout the indentation process. A 3-by-5 array of indentation marks with 50 µm spacing was made in each sample and the indentation force range was chosen to be 200–600 μN. This force range translated to an average depth of 5000 nm across multiple samples. The Oliver–Pharr method [24] was used in order to extract an elastic modulus from the load–displacement data.

### 2.7. Scanning Electron Microscopy (SEM) structural images

AC hydrogel samples with variations of the two components were prepared in 12-well plates following the same process described in Section 2.2. The samples were then fixed using a 4% paraformaldehyde solution (Electron Microscopy Sciences, Hatfield, PA, USA), and allowed to sit for 20 minutes. The samples were then cut into 10 mm by 10 mm squares, pre-frozen in –20 °C for an hour and then lyophilized at a vacuum pressure of 0.015 mBar and collector temperature of –52 °C for 3 days using a benchtop lyophilizer (Labconco, Kansas city, MO, USA). The dried samples were then imaged using a Quanta FEG scanning electron microscope (FEI) at 2–5 keV and 10–22 nA electron beam current.

### 2.8. Cell viability assay

A LIVE/DEAD^®^ cell viability kit (Molecular Probes L-3224), whose operation is based on the integrity of the plasma membrane, was utilized in accordance with the manufacturer’s protocol. AC hydrogels with variations in both components were synthesized and cells were kept in culture within these hydrogel environments for a week, while the media was changed every two days. The viability of cells was evaluated after this seven-day culture period. The dead cells were stained with ethidium homodimer (EthD-1, red stain) and the live cells were stained with calcein AM (CAM, green stain). The samples were then imaged using an Axio Observer D1 fluorescence microscope (Carl Zeiss AG) at an excitation wavelength of ~495 nm and the emitted wavelengths were filtered for 495 nm (CAM) and 635 nm (EthD-1). The images were then analyzed using the NIH ImageJ software in order to obtain the cell count for live/dead cells, and viability was calculated as the percentage ratio of live cells to the total number of cells. All viability experiments were repeated for four samples at each ECM composition.

### 2.9. Immunofluorescence staining and imaging

Alexa Fluor^®^ 488 phalloidin (Molecular Probes A12379) was used to visualize the filamentous actin structures of the cells in our 2D and 3D cultures. DAPI (Molecular Probes D1306) was also used to counterstain the nuclei of the cells. Briefly, the culture media was gently removed from the cellular constructs, and the constructs were carefully washed with DPBS. Cells were fixed using a 4% solution of paraformaldehyde for 20 minutes after which the solution was removed and the samples were thoroughly washed. The samples were then permeabilized with a diluted solution of 0.5% Triton X-100 (Sigma-Aldrich) for 15 minutes. The phalloidin solution with a 10 µM concentration was then applied and the samples were incubated for an hour in a dark environment. The phalloidin was then removed and samples were washed with DPBS. Finally, the DAPI solution with a 5 µM concentration was added and the samples were incubated for five minutes. The samples were imaged using the aforementioned microscope at an excitation/emission wavelength of 495/518 nm for the phalloidin and 358/461 nm for DAPI.

## 3. Results and discussion

### 3.1. Stiffness and porosity of hydrogel constructs

Whether in the case of a 3D matrix surrounding the cells omnidirectionally, or a 2D surface supporting sheets of cells, ECM stiffness is one of the few well-known mechanical cues, with significant influence on the biological performance of cellular constructs. One of the main advantages of composite hydrogels is the controllability of the gel stiffness and porosity by modulating the concentration of the hydrogel components in the pre-gel mixture [20]. We started by investigating the mechanical behavior of the agarose hydrogel without any additional components. A wide range of agarose hydrogels, all of which fall in the low polymer concentration domain of known gels, were studied. The concentration of agarose was increased from 0.5% to 3.0% (w/v) and the elastic modulus of the gel was observed to increase nonlinearly from 1.93 kPa to 7.55 kPa (Fig 2a). The agarose concentration was then fixed at 1% (w/v) and the collagen content of the hydrogel was increased from 0.05% to 0.2% (w/v). The addition of this secondary component increased the stiffness of the AC hydrogel, from 2.3 kPa up to a maximum of 10.53 kPa in the case of 0.2% collagen (Fig 2b). For the purpose of this study, we focus on neural tissue and the endothelium, both of which require extremely low stiffness values in the range of 1–10 kPa [25]. The elastic modulus values reported here cover this range of biologically relevant stiffnesses and this indeed demonstrates the suitability of the AC hydrogel to mimic the ECM from a mechanical perspective for both culture modes. Moreover, we have demonstrated that the elastic modulus could be easily configured within this range by modulating the hydrogel composition.

The lower stiffness observed in the 3% agarose hydrogel in comparison with the 1% agarose 0.2% collagen hydrogel with much less overall polymer content (1.2% w/v), indicates a distinction in the means by which agarose and collagen modulate the hydrogel stiffness. To further investigate such structural dissimilarities and gain better insight into the interactions between the components of our binary AC hydrogel, we imaged the porous structures of samples with variations in both components, using SEM. The agarose structures with very low collagen concentration (Fig 2c) resemble a mesh of intertwined polymeric nets. The mesh density is seen to increase with agarose concentration, resulting in a decrease in the average pore size. The addition of collagen above approximately 0.05% w/v (Fig 2d), meanwhile, results in the appearance of a more ‘wireframe’-like network in which thicker bundles of fibers appear generally near the edges of pores and have diameters as large as a few micrometers. Membranes of further ECM material now give the appearance of being ‘stretched’ between these bundles. Whether the ‘wireframe’ bundles are composed purely of the additional collagen or are a mixture of the constituent materials cannot be discerned from SEM images, but these bundles seem to act as a ‘backbone’ giving the polymeric network further structural support.

**Figure 2.**
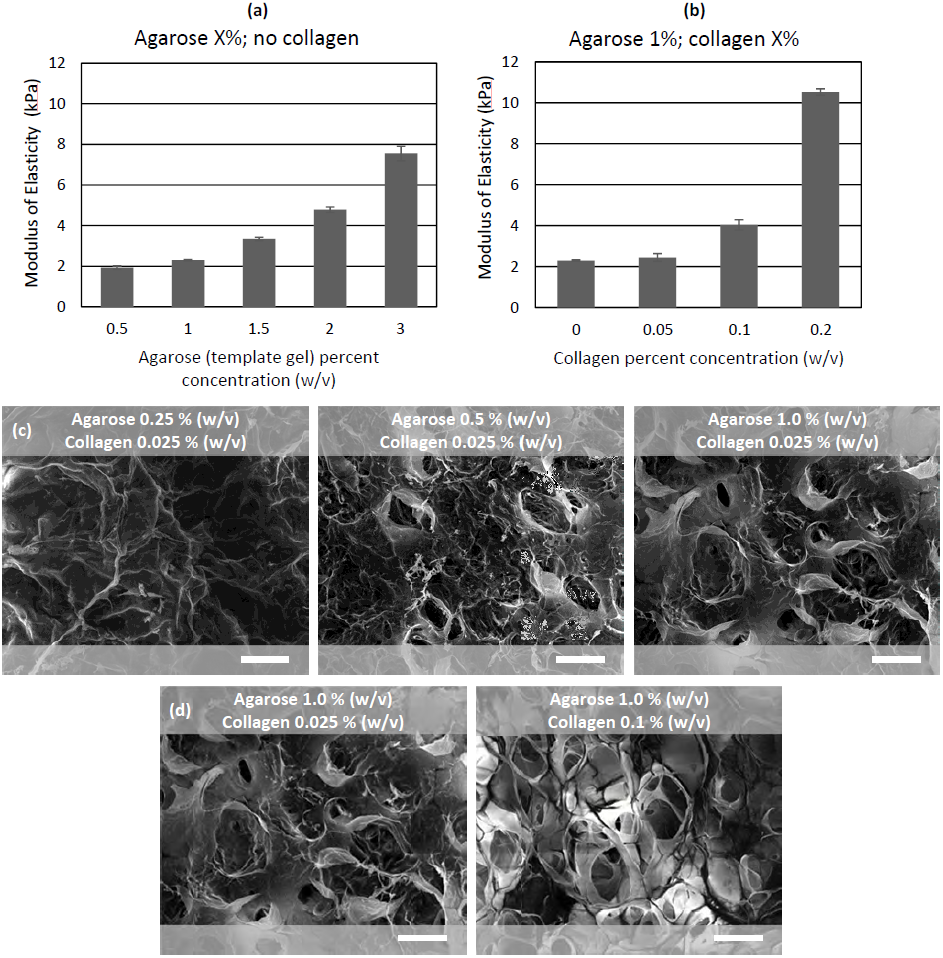
Mechanical properties of AC hydrogels and SEM images of gel structure; the effect of varying (a) ag arose concentration, and (b) co llagen concentration on hydrogel elastic m odulus. Stiffn ess values mat ch those of in-vivo cerebral and vascular tissue. SEM ima ges, indicating (c) a reduction in the avera ge diameter of pores at highe r agarose concentrations, and (d) a dded structur al support from fiber bundles highlighted by the presence of a thick ‘wireframe’. Scale bars are all 100 µ m.

There is clearly a distinction between the polymerization physics of the agarose and collagen components, and this could possibly explain the mechanical differences. Whatever the formation kinetics of these structures are, it is evident that the relationship between the agarose and collagen components is synergistic, since, as we observed in preliminary tests, collagen-only hydrogels — even at polymeric contents as high as 10 mg/mL — do not display adequate structural integrity, and are not suitable for a molding process.

Collagen is among the natural ECM molecules that allows for integrin binding of cells, due to the presence of RGD sequences that are exposed through denaturation and cleavage [26]. Therefore, these collagen bundles also serve as a binding target for the focal adhesion sites of the cells embedded in the AC hydrogel. This property will in turn directly affect cell proliferation and growth within the gel. However, as the collagen concentration increases, the entanglement of collagen fibers into larger fibers may result in lower surface-to-volume ratios. Hence, as we shall see in the following sections, the biocompatibility trend is not necessarily monotonically dependent on collagen content.

### 3.2. Fabrication results

AC hydrogels exhibit exceptional structural integrity, even at extremely low polymer concentrations and stiffness values as small as a few kPa, as shown in the previous section. This property is critical to the suitability of the hydrogels for the proposed method, as it allows us to transfer a 3D pattern accurately from the mold to the hydrogel. Therefore, we believe that composites of agarose hold great promise in fabricating 3D tissue constructs. Another important characteristic of agarose-based hydrogels is the significant hysteresis in melting and setting temperatures. We find this interesting property especially useful from a fabrication point of view as it directly translates to the high controllability of gelation speed and the exceptional stability of our constructs at operational temperatures.

Semi-cylindrical molds with diameters as small as 200 µm and acceptable surface roughness (approximately 5 μm r.m.s.) were fabricated using an inkjet 3D printer as described in the methods. These molds were used for the multilayer micro-molding process, enabling us to fabricate hydrogel constructs with vessel diameters ranging from 250 to 2000 µm (Fig 3a). Furthermore, coaxial structures with multi-layered walls of layer thicknesses as small as 50 to 100 µm were achieved (Fig 3b–d). Coaxial geometries with such fine feature sizes are beyond the capabilities of most bioprinting techniques [17]. The triple-layer constructs of Fig 3b–d show such 3D patterning capability, having an inner shell and outer region populated with SY5Y glial cells, and an intermediate layer without cells.

It can be observed from Figure 3 that the molded geometries are not perfectly circular in cross-section. We can identify two possible reasons for these geometrical deviations. Firstly, during production of the mold, the surface tension of the printed droplets of photocurable resin may cause rounding of the corners at the bases of the mold protrusions prior to curing. Secondly, after gelation and removal from the mold, the hydrogel’s own surface tension may lead to rounding of sharp corners. This second behavior is in fact potentially advantageous since it may serve to smoothen out any surface roughness transferred from the 3D-printed molds.

**Figure 3.**
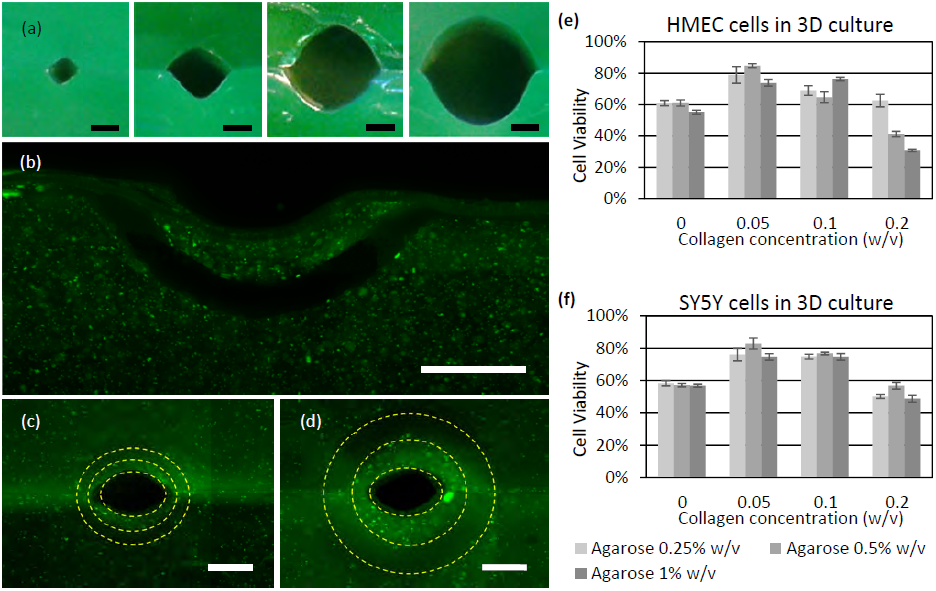
Fabricated constructs: (a) cross-sections of vessels with d iameters vary ing from 250– 1500 µm; (b) tri-layered construct with an inner diameter of 500 µm and layer thickness of 250 μm; enclosed channels with the same in ner diameter, and layer thick ness of (c) 125 μm, and (d) 2 50 μm; viabil ity analysis re sults for (e) SY5Y cells, and (f) HMEC cells, in 3 culture after seven days in vitro. Scale bars are all 500 μm. Error bars show standard deviation a mong four trials. Green stain in (b) –(d) shows live cells; in (a) it is simply mix ed into the gel.

### 3.3. Biocompatibility of hydrogel constructs

Cell viability, as a metric for the compatibility of the 3D microesnvironment with ce ll growth and proliferation, was assessed for all three cell types after seven days *in vitro*. Our results revealed a very interesting trend with respect to variations in either of the two components of the AC hydrogel. In the absence of collagen, the agarose hydrogel exhibits relatively poor viability (less th an 60%) across both cell types. Increasing the collagen content to an intermediate concentration (w/v) of 0.05–0.1% maximizes cell viability for both cell lines (Fig 3e–f). This result indicates that the presence of collagen in our composite hydrogel has an essential role in ensuring acceptable viability and enh anced biocompatibility of our constructs. Further increasing the collagen concentration reduces cell viability. The same trend can be seen for agarose: in most cases the composite with intermediate agarose concentration shows slightly better viability than the one with minimum concentration, and we observe a significant decrease in viability at very high concentration values.

These observations may be explained by the fact that although softer microenvironments are in favor of cell mobility and proliferation freedom, as suggested in the literature [22], there are mechanosensitive pathways such as MAPK that are found to be influential on cell proliferation and differentiation, the activation of which highly depends on the force transmitted between the interior of the nucleus, the cytoskeleton and the surrounding ECM [27], [28]. The focal adhesions are the intermediate links through which such signaling pathways between the ECM, the cytoskeletal actin structure and the nuclei are activated and such mechanical crosstalk may be disrupted in extremely low ECM stiffness values or in the absence of cell adhesive protein sequences in the ECM. Overall, the results indicate an optimal concentration of 0.5% w/v agarose and 0.05% w/v collagen at which the viability values for the SY5Y and HMEC cell lines were 82.8% and 84.7% respectively. These values indeed reveal the suitability of the AC hydrogel for 3D culture of both cell lines.

### 3.4. 3D growth and ECM remodeling

The cell proliferation, colonization and growth within the 3D AC hydrogel matrices were monitored at different hydrogel compositions. The SY5Y cells grown in 3D culture show very interesting dependences of both colony size and neurite extension on the hydrogel composition. At higher agarose concentrations, cells tend to become more isolated with less growth and aggregation, and fewer actin structures appear (Fig 4a–c). Moreover, at the lower agarose concentrations the 3D clusters of cells tend to be larger (Fig 4d–f). Additionally, neurite extensions are longer at lower agarose concentrations (Fig 4g–i). The effects of agarose concentration on cluster size and neurite extension can be explained by the data that we gathered from the mechanical characterization. We demonstrated that the stiffness of the hydrogel has a direct relationship with the agarose concentration whereas the pore size has an inverse relationship. Since the stiffer hydrogels show less compliance and deformation, the prolifertion freedom and mobility are also correspondingly limited. The ability of the cells to attach themselves to the agarose backbone and extend their neurite protrusions into the hydrogel, as well as their extension range, is determined by their ability to penetrate the hydrogel, which is inversely correlated with the hydrogel stiffness.

**Figure 4.**
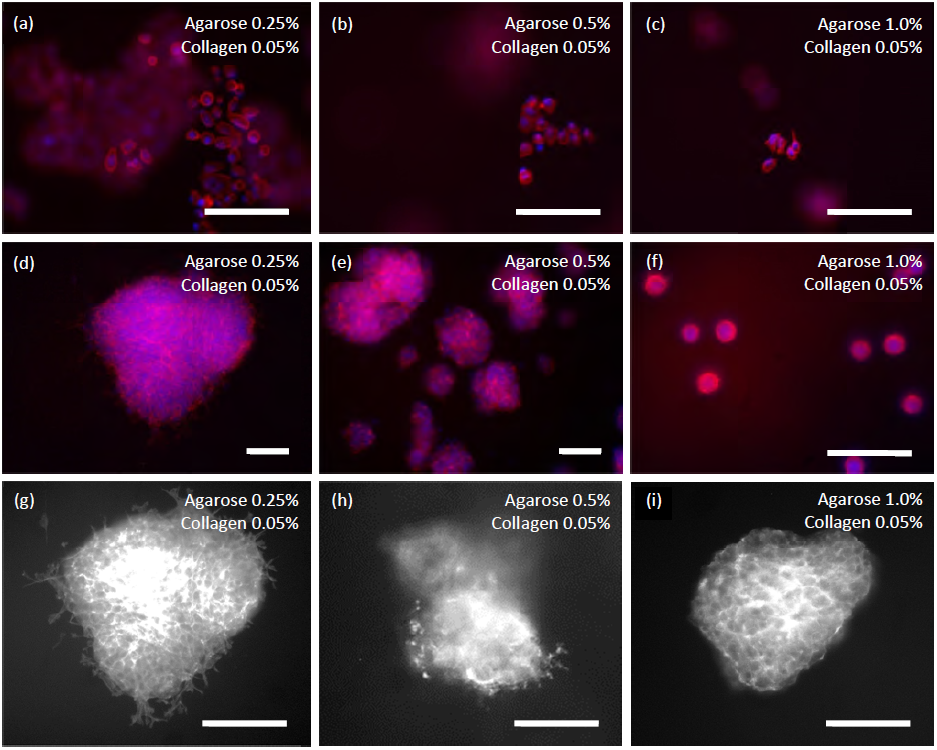
Expansion and colonization o f SY5Y glial c ells within the 3D AC hydrog el: (a–c) overall trend of cell population reduction with the increase in agarose co ncentration fro m 0.25% observed at different locations ac ross the samples (a) to 1% (c); (d–f) decrease i n colony size of cellular clusters with the increase of agar ose concentration from 0.25% (d) to 1% (f); (g–i) decrease in length of neuri te extensions w ith the increa se of agarose c oncentration from 0.25% (g) to 1% (i). Scale bars are all 50 μm. Stainin g: a–f: red: actin, blue: nucl ei; g–i: actin.

### 3.5. 2D culture and lumen formation

In order to model a vascular network successfully, we have to achieve exceptional surface adhesion properties to facilitate lumen formation. Adhesion of the cells to the hydrogel substrate is an essential precursor to inter-cellular tight-junction formation. The AC hydrogels, however, lack such natural cell-adhesive properties and therefore we experimented with a variety of surface coatings that could potentially improve the cell-adhesive properties of our substrate. We have focused on modeling blood vessels located in the brain, which have distinctive characteristics. These blood vessels have a multi-luminal structure composed of a thin endothelium surrounded by glial cells. We have studied the spreading behavior of endothelial and glial cells, each of which plays a special role in the integrity of this neurovascular unit. Single-layer coatings of Matrigel, human fibronectin, collagen and PDL were used for the endothelial cells. Matrigel and PDL coatings were used for the glial cells. Moreover, the sequential application of multiple coatings was studied for several of the coating combinations, inspired by the work of Han *et al.* [29]. For this purpose, a 0.25% w/v PDL solution was used to produce the first coating layer that would facilitate the adhesion of a secondary coating of Matrigel, collagen or a combination of both.

Considering first the effect of the substrate configuration, at higher agarose concentrations HMEC spreading percentage is significantly increased (Fig 5g), showing that higher stiffness values enable better 2D spreading of cells — as would be expected considering that plastic culture surfaces perform exceptionally well for 2D cell culture. Here, spreading percentage is defined as the fraction of cells that exhibit a non-spherical shape and/or one or more filopodia. Moreover, decreasing the thickness of the agarose layer — which is effectively increasing the stiffness of the surface because the underlying container is much stiffer than the agarose — also leads to better cell-spreading properties (Fig 5g).

For the hFN and collagen coatings, the cells do not attach to the AC hydrogel surface at all. However, the HMEC cells easily adhere to hydrogel surfaces with PDL and Matrigel coatings. Although the cell-spreading percentages of the HMEC cells on the AC hydrogel with only the PDL coating are very low (11.2%), spreading is considerably improved in the presence of thin layers of Matrigel (81.3%) and collagen (89.9%) (Fig 5h).

Turning to cell-spreading on coating combinations, we observe significant enhancements in surface coverage and confluence over single-material coatings. These coatings also show acceptable cell spreading (above 50%, Fig 5h) despite the fact that cells still do not form a fully confluent monolayer within the first two days. Overall, the collagen-only coating results in the highest percentage of spread cells; however, the surface coverage of this coating within the same two-day period is not as good as the multi-material coatings. It is noteworthy that we tried multiple solutions of collagen, Matrigel and PDL in order to find the optimum concentrations, which were 5 mg/mL, 200 μg/mL and 0.25 mg/mL respectively.

The same study was carried out with SY5Y glial cells, and we realized that the only coatings that facilitated SY5Y attachment were collagen and PDL. The PDL concentration range that gave optimal cell adhesion was found to be 0.1% to 0.25% w/v (the 0.1% w/v case is shown in Fig 5a–b). One of the issues that we encountered was the separation of the layer of cells from the substrate after the second day of culture. This instability could be attributed to the over-confluence of the layer and breakage of the adhesion complexes as a result of layer expansion. The addition of a collagen coating gave better stability to the layer of cells in spite of the over-confluence issue (Fig 5c).

**Figure 5.**
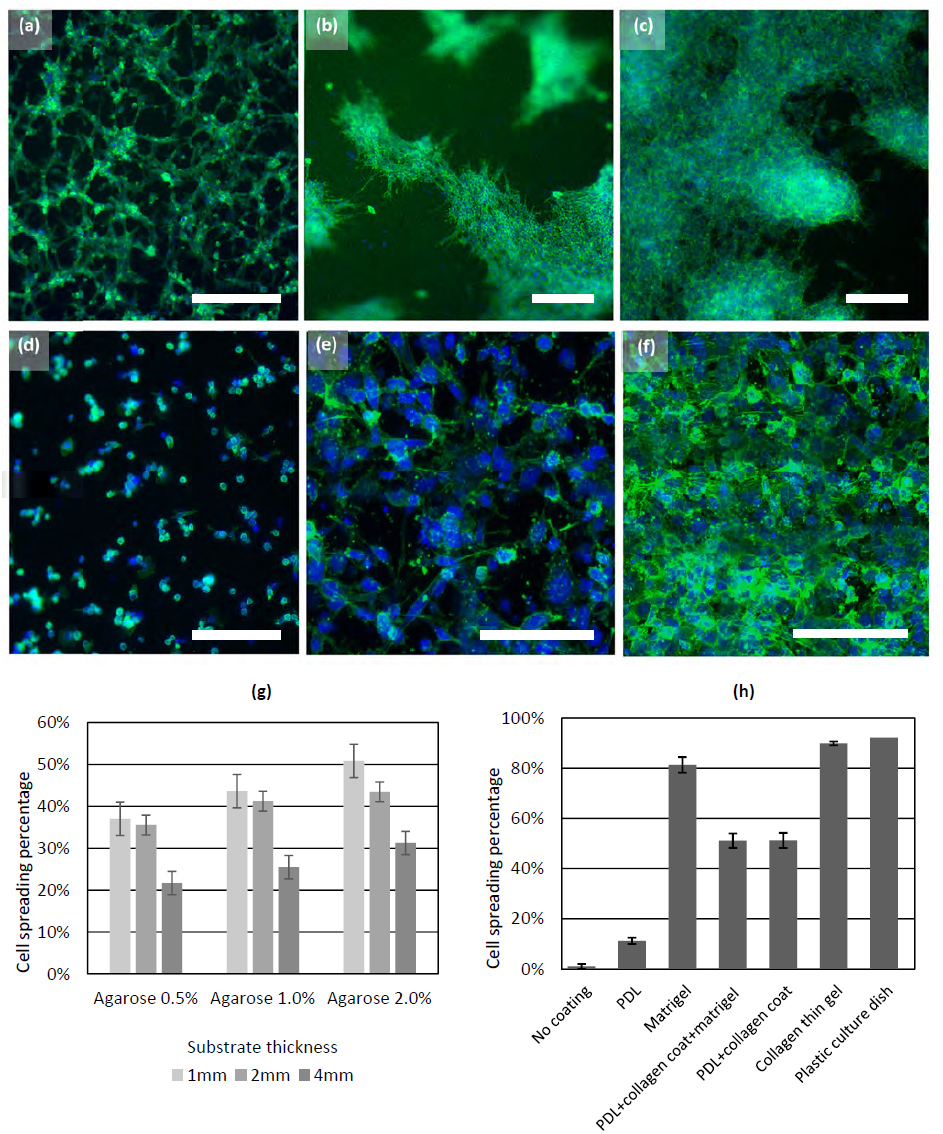
Modulating the surface adhe sion properties of the AC hyd rogel for SY5 Y and HMEC c ells. Adhesion of SY5Y cells on th e optimal coat ing (0.1% w/v PDL) and sub strate (2 mm-thick layer of 0 .5% agarose a nd 0.05% coll agen): (a) cell adher ence after one day in vitro; ( b) accumulation of cells on the surface after two days in v itro; (c) presence of a collagen coating on top of the PDL ena bling stable at tachment of th e layer of cells even after th ree days. Adhesion of HMEC cells on different coatings: (d) PDL coating found to be unsuitable for HMEC cells; (e) best cell spreading achieved using a 100 μm-thick layer of collagen gel; (f) the same configuration as (e); however, the addition of a sublayer of 3D cultured SY5Y cells underneath the endothelium substantially improves surface coverage and area confluence. Quantitative analysis of the spreading of HMEC cells on various substrate configurations: (g) modifications in the hydrogel substrate thickness and polymer content (collagen concentration is zero, and the agarose concentration is modulated), and (h) application of surface coatings (again, the substrate is a 2 mm-thick layer of 0.5% agarose and 0.05% collagen). Scale bars are all 100 μm. All images: green stain: actin, blue stain: nuclei.

### 3.6. Multi-layered 3D vascular constructs

Using the optimal configurations that were obtained from the previous sections, a triple-layer construct was fabricated. The outermost layer of the construct, with an inner diameter of 1 mm, was made of a 0.5% agarose, 0.5% collagen AC composite gel, and was populated with SY5Y cells. A thin intermediate layer of 0.5% collagen gel with an inner diameter of 0.5 mm seeded with SY5Y cells was then cast and finally the endothelial cells were seeded on the inner surface of this layer. The measurements and imaging of the construct were carried out after three to four days *in vitro*. Prior to this 3D experiment, the effect of the presence of glial cells on the confluence and area coverage of the endothelia was investigated in the case of a 2D stack and our results indicated a significant improvement from the endothelium-only case (Fig 5e) to the co-culture case where the glial cells were also present in the layer underneath the endothelium (Fig 5f). The same observation was made in the 3D case. The overall triple-layered lumen and its projections from different angles are shown in Fig 6. The majority of the cells seen are the HMECs lining the lumen. The SY5Y glial cells in the intermediate cast layer can be seen scattered in the region around the lumen. The SY5Y cells in the outermost cast layer have a smaller density than those in the intermediate layer and hence appear very sparsely in the images.

**Figure 6.**
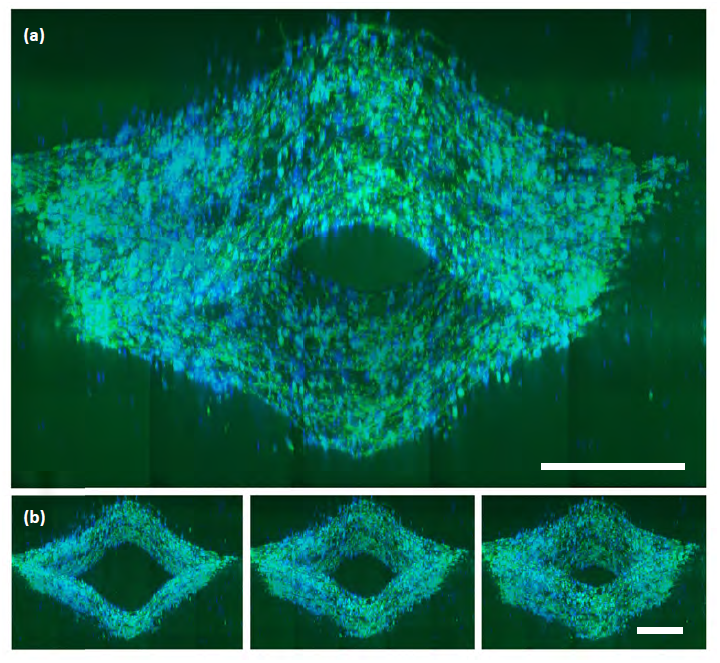
3D vascular co nstruct: (a) en dothelia with glial cells surrounding the lu men at a close proximity (b) projection s of the same l umen. Scale ba rs are 200 μm. Green stain: actin, blue sta in: nuclei.

## 4. Conclusions

We have introduced a versatile method to create multi-layered, cell-laden hydrogel microstructures with coaxial geometries and heterogeneous elastic moduli. The method could be tuned towards vasclar networks lying in the circulation interface with neural tissue. The main advantages of the presented fabrication method are: (1) scalability across multiple vessel diameters ranging from 200 to 2000 μm, and the posibility to shrink dimensions further by enhancing the mold fabrication technique, (2) ease of sample preparation and culture convenience, making the setup readily available to any tissue culture facility and eliminating complications encountered with microfluidic systems and bioprinters, and (3) the capability to produce micro-features as small as the mold preparation technology is capable of defining. This fabrication route enables the production of an *in-vitro* model with following advantages over state-of-art *in-vitro* models: (1) establishing a co-culture with multiple cell-types in **direct contact** with each-other, (2) incorporation of heterogeneous elastic modulus for **optimal growth** of different cell types, and (3) controllability of **thickness, cellular density and composition** of each tissue layer.

## 5. Acknowledgements

This work was supported by faculty start-up funding from the U.C. Berkeley Department of Mechanical Engineering. The authors gratefully acknowledge the assistance of the staff of the Biomolecular Nanotechnology Center, the Molecular and Cell Culture Facility, operated by Biosciences Divisional Services, and the Jacobs Institute for Design Innovation, all at U.C. Berkeley. In particular, valuable input from Alison Killilea, Paul Lum, Chris Parsell, Carissa Tasto, and Mary West is acknowledged. H.T. acknowledges helpful discussions with Roger Kamm and the authors acknowledge helpful discussions with Yasaman Daghighi.

